# Non-syndromic autism *SCN2A* variants selectively exert dominant-negative effects on Na_v_1.2 channels

**DOI:** 10.64898/2026.02.24.707547

**Authors:** Sandrine Cestèle, Renzo Guerrini, Sandra Difhallah, Davide Mei, Natalie Leroudier, Marialuisa Ricci, Simona Balestrini, Massimo Mantegazza

**Affiliations:** University Cote d’Azur, 06560 Valbonne-Sophia Antipolis, France; CNRS UMR 7275, Institute of Molecular and Cellular Pharmacology (IPMC), 06560 Valbonne-Sophia Antipolis, France; Inserm U1323, 06560 Valbonne-Sophia Antipolis, France; Neuroscience Department, Meyer Children’s Hospital, 50139 Florence, Italy; University of Florence, 50139 Florence, Italy

**Keywords:** sodium channels, autism spectrum disorder, epilepsy, developmental epileptic encephalopathies, neurodevelopmental disorders, excitability

## Abstract

The voltage-gated Na^+^ channel Na_v_1.2 has a key role for the initiation and propagation of action potentials and therefore in neuronal excitability in brain development and function. Genetic variants of the encoding gene *SCN2A* cause various neurodevelopmental phenotypes with infantile-childhood onset. Here, we investigated the functional impact on hNa_v_1.2 function of 15 variants associated with pure non-syndromic ASD (nsASD), ASD with epileptic activity, developmental and epileptic encephalopathy or schizophrenia. Only nsASD variants caused a complete loss of function hNa_v_1.2 channels when expressed alone in tsA-201 cells and cultured neocortical neurons. Co-expression of the WT and mutant channels mimicking heterozygosis revealed that ASD mutants induce a dominant negative effect. Using different strategies to impair the domains of the channels that have been suggested to be implicated in the interaction of two Na_v_ α subunits, we reversed the dominant negative effect of ASD mutants on WT channels. These findings identify in heterologous systems a mechanistically distinct class of *SCN2A* variants implicated in nsASD, defined by dominant-negative loss of Nav1.2 function, with potential utility as a biomarker for genetic counseling, patient stratification, and the development of precision therapeutic strategies.

## INTRODUCTION

The *SCN2A* gene encodes the voltage-gated Na^+^ channel α subunit Na_v_1.2, which is the main Na^+^ channel responsible for the initiation and propagation of action potentials of cortical excitatory neurons in the early postnatal period, while at a later age it is also involved in their dendritic excitability and probably in setting features of synaptic transmission^1–3^. Heterozygous pathogenic variants of *SCN2A/*Na_v_1.2 can cause a wide phenotypic spectrum, including mild epilepsy, different types of developmental and epileptic encephalopathies (DEEs) or neurodevelopmental disorders without epilepsy^4,5^.

Some *SCN2A/*Na_v_1.2 variants cause phenotypes with late infantile or childhood onset, including infantile-childhood DEEs (ICDEEs) and different neurodevelopmental disorders^6,7^. ICDEE patients with onset between 3 months and 1 year of age often exhibit an infantile spasms syndrome phenotype, whereas ICDEE patients with onset after 1 year of age may have variable epilepsy phenotypes that cannot be classified within a specific epilepsy syndrome, developmental delay/intellectual disability and autistic traits. *SCN2A/*Na_v_1.2 neurodevelopmental disorders include autism spectrum disorder (ASD) and/or intellectual disability, as well as other neuropsychiatric conditions such as schizophrenia^2,6,7^. Although seizures have been observed in some of these patients after signs of neurodevelopmental dysfunctions became apparent, epilepsy is not a major feature of their phenotype. Large-scale human genetic studies have indicated that *SCN2A/*Na_v_1.2 variants are among the leading genetic causes of non-syndromic ASD (nsASD)^8,9^.

Other *SCN2A/*Na_v_1.2 variants cause phenotypes with onset in the first 3 months of life, mostly in the neonatal period^6,7^. Some of these patients exhibit self-limited neonatal/infantile epilepsy^10^, others exhibit severe neonatal-early infantile DEE (NEIDEE) phenotypes with drug-resistant seizures and intellectual disability.

To disclose pathological mechanisms, stratify patients and identify specific therapies in a precision medicine framework, it is important to identify the functional effects of the variants and correlate them to the phenotype. Functional studies in transfected cell lines have provided some genotype-phenotype correlations, showing that gain-of-function (GoF) variants cause self-limited epilepsy or NEIDEE, which can respond to treatment with Na^+^ channel blockers, whereas loss-of-function (LoF) variants cause ICDEE or neurodevelopmental disorders without epilepsy, phenotypes that are generally worsened by Na^+^ channel blockers^1,2,7^. However, it has been thus far more difficult to identify clear genotype-phenotype relationships within GoF or LoF variants^1,5,11,12^.

It has been proposed that *SCN2A/*Na_v_1.2 ASD variants, compared to ICDEE ones, cause larger LoF that lead to haploinsufficiency^13^. Haploinsufficient *Scn2a*^+/-^ mice, especially at young age, show autistic-like features^3,14–17^, which are milder than those observed in mice with a larger reduction of *Scn2a* expression^18–21^. Thus, pathological mechanisms that can cause LoF larger than haploinsufficiency could be involved in *SCN2A/*Na_v_1.2 ASD. Although all these variants have been identified in heterozygous patients, functional studies have been performed thus far with conditions that mimic homozygosis. However, it has been proposed that α subunits, including Na_v_1.2, can interact and form dimers^22^. This interaction may generate dominant negative effects when wild-type and mutant proteins are co-expressed, as in conditions of heterozygosis, in which the mutant subunit may reduce the function of the wild-type one^23^.

Here we performed functional studies in transfected cell lines and neurons of *SCN2A/*Na_v_1.2 variants that cause phenotypes with late infantile or childhood onset, in which we reproduced the conditions of heterozygosis. We investigated published variants and novel variants identified in a newly described cohort of patients. We identified dominant negative effects as a novel specific pathological mechanism for variants involved in nsASD.

## MATERIALS AND METHODS

### Plasmid and Mutagenesis

We used the cDNA of the human Na_v_1.2 channel α subunit (hNa_v_1.2, GenBank accession no. NM_021007) provided by Dr. Jeff Clare (Glaxo-SmithKline, Stevenage, Herts, United Kingdom), which we subcloned into the pCDM8 vector to minimize rearrangements^24^. The cDNA of the human Kir4.1 channel (GenBank accession no. NM_002241) was obtained from OriGene technologies Inc., USA (CAT#: SC118741). We introduced the mutations with the Quick-Change Lightning Kit (Stratagene) as already described^25,26^; primers’ sequences are available on request. To isolate the Na^+^ currents generated by the transfected channels from the endogenous ones in the experiments in which we used neocortical neurons, we expressed hNa_v_1.2 channels (WT or carrying a pathogenic mutation) resistant to the specific blocker tetrodotoxin (TTX), in which the phenylalanine at position 387 was replaced with a serine^24–26^. The plasmid containing difopein (dimeric fourteen-three-three peptide inhibitor)^27^ fused to YFP (pEYFP-C1-difopein) was provided by Dr. Isabelle Deschênes (Case Western Reserve University, Cleveland, USA)^22^.

### Cell culture and transfection

We used the cell line tsA-201, maintained and transiently transfected with CaPO_4_ as already reported^25,26,28^. Neocortical neurons were prepared from E17 mouse embryos (Charles River) and maintained in primary culture as already described^25,29^. Transfections of neurons were performed with Lipofectamin 2000 (Invitrogen) 5 days after the preparation and recorded 24-48h after the transfection. We co-transfected a plasmid expressing Yellow Fluorescent Protein (pEYFP-N1; Clontech) to identify the transfected cells (tsA-cells 201 or neocortical neurons) for electrophysiological recordings. For recordings from cultured neurons, we selected cells with pyramidal morphology^30^.

### Electrophysiological recordings and analysis

We used the whole-cell configuration of the patch-clamp technique to record Na^+^ currents, as previously described^25^. Recordings were realized at room temperature (20-24°C) with a Multiclamp 700A amplifier and pClamp 10.2 software (Axon Instruments/Molecular Devices). Signals were filtered at 10 kHz and sampled at 50 kHz. Electrode capacitance and series resistance were compensated during the experiments. Pipette resistance was between 2-2.5 MOhms and voltage error maintained under 2.5 mV. The P/4 subtraction paradigm was used to cancel the remaining transient and leakage currents. Recording solutions for tsA-201 cells were (in mM): external solution 150 NaCl, 1 MgCl_2_, 1.5 CaCl_2_, KCl_2_ and 10 HEPES (pH 7.4 with NaOH); internal pipette solution 105 CsF, 35 NaCl, 10 EGTA, 10 HEPES and (pH 7.4 with CsOH). Recording solutions for neurons were (in mM): external solution 140 NaCl, 2 MgCl_2_, 2 CaCl_2_, 1 BaCl_2_, 1 CdCl_2_, 10 HEPES (pH 7.4 with NaOH) and TTX 1 µM; internal pipette solution (in mM): 130 CsF, 10 NaCl, 10 EGTA and 10 HEPES (pH 7.4 with CsOH). The recordings were started 5 minutes after reaching the whole-cell configuration to allow a complete dialysis of the cytoplasm. Voltage dependence of activation was studied applying test pulses of 100-ms from −110 mV to +60 mV from a holding potential at −120 mV. Voltage dependence of inactivation was studied with a 100-ms prepulse at different potentials followed by a test pulse at −10 mV. Conductance-voltage curves were derived from current-voltage (I–V) curves according to G = I/(V−Vr), where I is the peak current, V is the test voltage, and Vr is the apparent observed reversal potential. The voltage dependence of activation and the voltage dependence of inactivation were fit to Boltzmann relationships in the form y = 1/(1 + exp((V1 2 −V)/k)), where y is normalized GNa or INa, V_1/2_ is the voltage of half-maximal activation (Va) or inactivation (Vh) and k is a slope factor. Action potential clamp recordings were performed as already described^26,31–33^. The inter-sweep interval was 8s for all the protocols. Data were analyzed with pClamp v10.2 (Axon Instruments/Molecular Devices) and Origin2021 (OriginLab). Junction potential was not corrected.

### Binding assay

The toxin AaHII, isolated from the venom of the scorpion *Androctonus australis Hector* was a generous gift of Dr. MF Eauclaire and Dr. P. Bougis. It was radioiodinated and used for binding experiments as described in^34,35^. We selected this site–3 toxin because it yields clean, highly sensitive binding, although some mutants could not be assessed, as they lack its binding site^36^. Binding experiments were performed on intact transfected cells in binding buffer solution (Choline Chloride 130 mM, HEPES 50 mM, Glucose 5.5 mM, MgSO40.8 mM, KCl 5.4 mM, BSA 1 mg/ml, pH = 7.4) with the addition of 10 mg/ml gramicidin A for generating a more negative membrane potential. Cells were incubated in the presence of 125I-AaHII at 37 °C for 45 min and unbound toxin was removed by aspirating each well and rinsing 2 times using the washing buffer (Choline Chloride 163 mM, HEPES 5 mM, CaCl2 1.8 mM, MgSO4 0.8 mM, pH = 7.4). Adherent cells with bound toxin were dissolved in 1 ml of 0.4 N NaOH per well and total cell protein was determined using bovine serum albumin as a standard in a modified Lowry protein assay (Bio-Rad Protein Assay).

### Clinical study

Clinical assessment comprised standardized neurological, neuropsychological, and neuropsychiatric evaluations. Patients were followed longitudinally for a median duration of 10 years (range, 4–14 years). Available brain MRI scans and EEG recordings obtained during wakefulness and sleep were systematically reviewed, with particular attention to background activity, physiological patterns and epileptiform discharges. Epilepsy syndromes were classified according to the International League Against Epilepsy (ILAE) criteria. Treatment efficacy was categorized based on longitudinal clinical records and the treating physician’s assessment. Neuropsychological and neuropsychiatric features were evaluated using validated instruments when feasible, including the Autism Diagnostic Observation Schedule–Second Edition (ADOS-2), Autism Diagnostic Interview–Revised (ADI-R), Vineland Adaptive Behavior Scales, and Leiter International Performance Scale–Revised (Leiter-R). When formal testing was not possible, assessments were based on structured clinical observation focusing on communication abilities, adaptive functioning, and behavioral features. All clinical data were entered into a pseudonymized database and independently verified for accuracy. Phenotypic classification was determined based on neurological history, EEG findings, and developmental trajectory by investigators blinded to the functional data. Clinical and functional data were subsequently analyzed jointly. For each individual, genotype–phenotype correlations were explored by comparing in vitro functional data with the clinical presentation, with the aim of identifying potential associations between the functional mechanism of the variant (dominant-negative versus loss-of-function) and the observed phenotype. The study was approved by the Paediatric Ethics Committee of Meyer Children’s Hospital IRCCS (DECODEE project, Florence, Italy). Written informed consent for genetic testing and for the use of anonymized clinical data for research purposes was obtained from all participants’ parents or legal guardians.

### Statistical analysis

Datasets were tested for normal distribution with the Kolmogorov–Smirnov test; and homogeneity of variance with the Brown-Forsythe test. Groups with normal or Lognormal distribution were compared with Student’s t-test or one-way ANOVA followed by Dunnet’s T3 post hoc test. Groups not normally distributed or with n too small to test for normality were compared with the Kruskal-Wallis test followed by Mann-Whitney post-hoc test with Bonferroni correction for multiple comparisons. *p* < 0.05 was considered significant. Significance is indicated in the figures as: *p < 0.05, **p < 0.01, ***p < 0.001, ****p < 0.0001. If not differently indicated, data are shown as means ± SEM, ‘‘n’’ indicates the number of cells. Statistical tests were performed with Prism 10.6 (GraphPad) or Origin 2025 (OriginLab).

## RESULTS

### Clinical features of patients carrying SCN2A/Nav1.2 variants investigated in this study

We performed the functional study of 15 *SCN2A*/Na_v_1.2 variants identified in patients with different phenotypes (Fig.1). Numerous variants are part of a cohort of patients of the Meyer Children’s Hospital IRCCS (Florence, Italy), which includes variants that were not reported before (Supplementary Table 1). Some other variants that we studied were selected from the literature: R379H, R937H, C959X and G1013X had been identified in patients with nsASD^9,37^; R850P and V1289F had been identified in patients with schizophrenia^38^; T1420M had been reported in a patient classified as nsASD, with no further detail available in the original publication^39^. However, based on the results of our functional analysis presented here, we can hypothesise a close similarity of the phenotype of the T1420M patient with that observed for the patient of the Meyer’s cohort carrying the R1635Q variant. We selected the medical records of patients carrying the L1314P and R1635Q variants for a detailed report. Additional information about these two patients and others of the Meyer’s cohort are reported in Supplementary Table 1.

**Figure 1:**
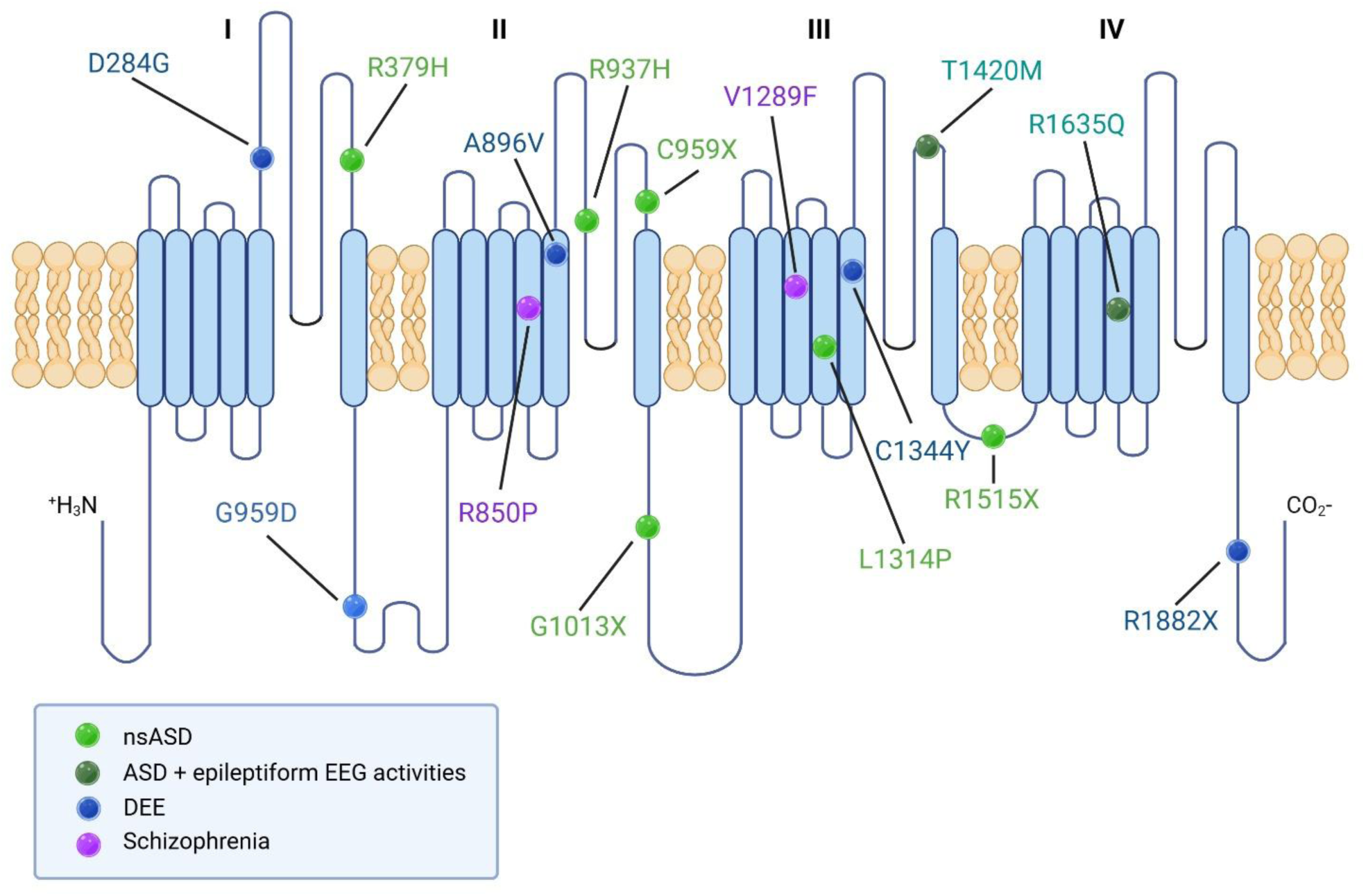
Localization on the hNa_v_1.2 protein of the 15 genetic variants that we have characterized. The α subunit consists of four homologous domains (I-IV) containing 6 transmembrane segments (S1-S6) connected by extra- and intra-cellular loops. The voltage sensor is present in the S4 segments of each domain and the pore region is formed by S5-S6 segments. nsASD: non-syndromic Autism Spectrum Disorder; DEE: Developmental and Epileptic Encephalopathy.

The patient carrying the L1314P variant was a 15-years-old girl, brought to neurological attention at age one year due to global developmental delay and microcephaly. Brain MRI performed at 18 months was unrevealing, except for the small brain size. Neuropsychological assessment at age 5 years revealed severe intellectual disability (ID) and absent speech. Since age 3 years the patient had manifested atypical attitudes in social interaction and communication skills, evaluated with ADOS-2 test (module 1), that revealed severe autism spectrum disorder. No clinical seizures were observed or reported. The only EEG abnormality observed was an increased amplitude of vertex sharp waves during the wakefulness-to-sleep transition (Supplementary Fig.1). At age of 14 years a non-verbal scale (Leiter-R) was consistent with moderate ID.

The patient carrying the R1635Q variant, a boy aged 12-years at the time of the study, was initially brought to medical attention at 3 years of age due to global developmental delay with absent speech and impaired communication skills. Brain MRI was normal. Subsequent evaluations highlighted autistic features with persistently absent speech, poor eye contact and social interaction, motor stereotypies, hypersensitivity to loud noises, no interest in interaction with peers. Although a training program with the applied behavior analysis (ABA) method was soon started with some progress, at age 6 years the ADOS-2 test (module 1) scores were consistent with severe autism spectrum disorder. Formal cognitive testing could not be administered. The child remained nonverbal and could only express his requests through vocalizations and gestures. EEG recordings showed continuous bilateral centro-temporal spikes (Supplementary Fig. 2), which were greatly activated during sleep; no clinical seizures were observed. The R1635Q variant was previously identified in patients with epilepsy and neurodevelopmental disorders, in which epilepsy is a defining feature^40^ (a phenotype aligning with the DEE spectrum), and in a patient described as autistic without observed clinical seizures^41^, suggesting phenotypic similarities with the patient carrying this variant in our cohort.

Other patients in the cohort of the Meyer Children’s Hospital IRCCS carried the D284G, A896V, G659D, C1344Y and R1882X variants, and had ICDEE phenotypes, in which different seizure types, with variable outcomes, were associate to mild to severe intellectual disability (ID) (Supplementary Table 1). Some of these patients exhibited syndromic autistic phenotypes within a DEE framework.

### ASD-hNa_v_1.2 mutants expressed in isolation in tsA-201 cells show complete/nearly complete LoF

To investigate genotype/phenotype relationships of *SCN2A* mutants, we introduced variants into the cDNA clone of the human Na_v_1.2 Na^+^ channel (hNa_v_1.2) by site directed-mutagenesis. Using patch-clamp whole-cell recordings on transiently transfected tsA-201 cells, we analyzed the functional properties of mutants comparing with those of WT hNa_v_1.2. We first describe the variants that, considering available clinical data and our functional findings, we consider as occurring in nsASD patients: R379H, R937H, C959X, G1013X, L1314P, R1515X (Fig.1).

Representative traces of Na^+^ current elicited with a series of depolarizing steps in tsA-201 cells transfected with WT or ASD mutant channels expressed alone are shown in Fig.2a. Fig.2b displays the quantification of the maximal current density (to normalize for the size of the cells) and indicates that five of the six ASD mutants tested display no current, whereas one, R379H, shows strongly reduced (80%) current density. These data indicate nearly complete LoF for the 6 ASD mutants tested. To better disclose the overall impact of the variant R379H on the function of hNa_v_1.2 in a dynamic condition, we applied as voltage command a neuronal discharge, as we did in previous studies^25,31,32^, and recorded the corresponding Na^+^ action currents quantified as current densities (Fig.2c). The comparison of the peak current density of the first action current and the mean peak current densities of the last 3 action currents (Fig.2d) indicates that the R379H induces 76% and 94% reduction, respectively, which results in a major LoF of hNa_v_1.2. Therefore, our data confirm previous results showing that nsASD *SCN2A* variants cause a severe LoF^13^.

**Figure 2:**
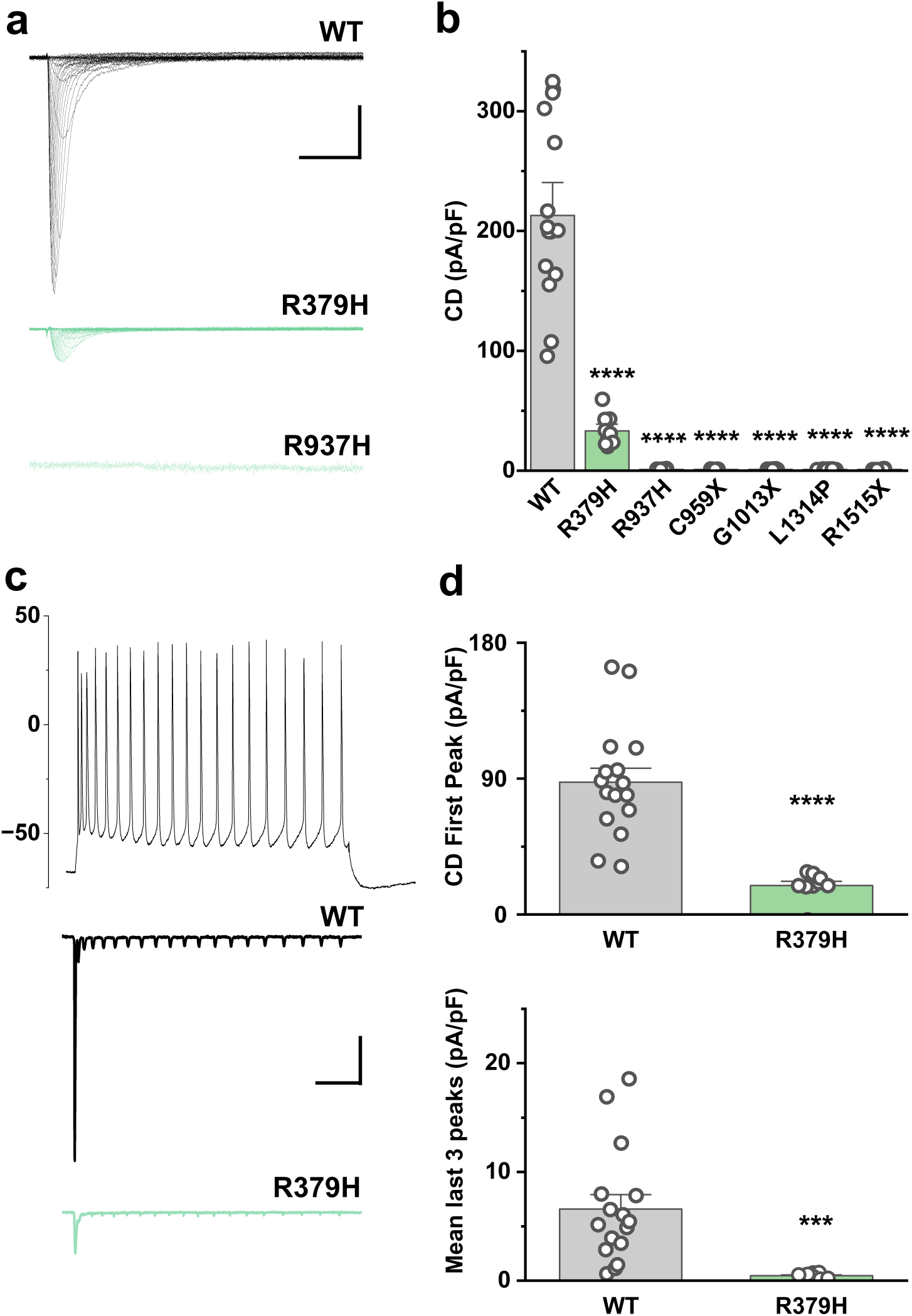
ASD-hNa_v_1.2 mutants expressed in tsA-201 cells show complete/nearly complete loss-of-function. **(a)** Representative whole-cell Na^+^ current families for hNa_v_1.2 WT and ASD mutants recorded with depolarizing voltage steps from -80 to +60 mV in 5 mV increments form a holding potential of -100 mV. Among the mutants with no current, only R937H is shown; scale bars 1nA, 4ms. **(b)** Maximal current density calculated for tsA-201 cells transfected with the WT or the ASD mutants. **(c)** Action Na^+^ currents recorded using a voltage stimulus an action potential discharge (first panel) for WT or R379H channels (data are expressed as mean of current density, error bars are not shown for clarity); scale bars 20pA/pF, 20ms. **(d)** Comparison of the current density of the first action current and of the mean of the last three action currents in the discharge. Data shown as mean±SEM. See Supplementary material (Statistical Tables) for values, n and statistical tests.

### ASD hNav1.2 mutants are not rescued in neocortical neurons and show dominant negative effects

To determine whether a neuronal cell background rescues plasma membrane targeting of hNav1.2 mutants, as we have previously shown for other Na^+^ channel mutants^25,29^, we expressed ASD-associated mutants in primary neocortical neurons. Using TTX-resistant (F385S) constructs in the presence of 1µM TTX to isolate exogenous currents^25,29^, we found that five mutants failed to generate measurable current, while R379H reduced current density by 69% (Fig.3a). These results indicate that the LoF observed in tsA-201 cells persists in a neuronal context, showing that the cellular background does not influence the functional deficit.

Then, to mimic the heterozygous condition observed in patients, we co-expressed WT and mutant hNa_v_1.2 channels in neocortical neurons (1:1 molar ratio, with 50% of the standard WT cDNA) (Figure 3b). As a control, to evaluate the effect of co-expression of a membrane protein that does not directly interact with hNa_v_1.2 but can compete with it for protein synthesis and trafficking, we co-transfected neurons with WT hNav1.2 and hKir4.1, a non-interacting K⁺ channel, which did not affect WT current density (Fig.3b). Strikingly, co-expression with any of the ASD mutants significantly reduced WT current density: R379H by 38%, L1314P by 52%, R937H and R1515X by 53%, and C959X and G1013X by 58% (Fig.3b). Notably, even R379H, which retained partial function when expressed alone (Fig.3A), showed a dominant-negative effect in the presence of WT channels. These data demonstrate that all the ASD-associated hNa_v_1.2 mutants we tested impair WT channel function, consistent with a dominant-negative mechanism.

**Figure 3:**
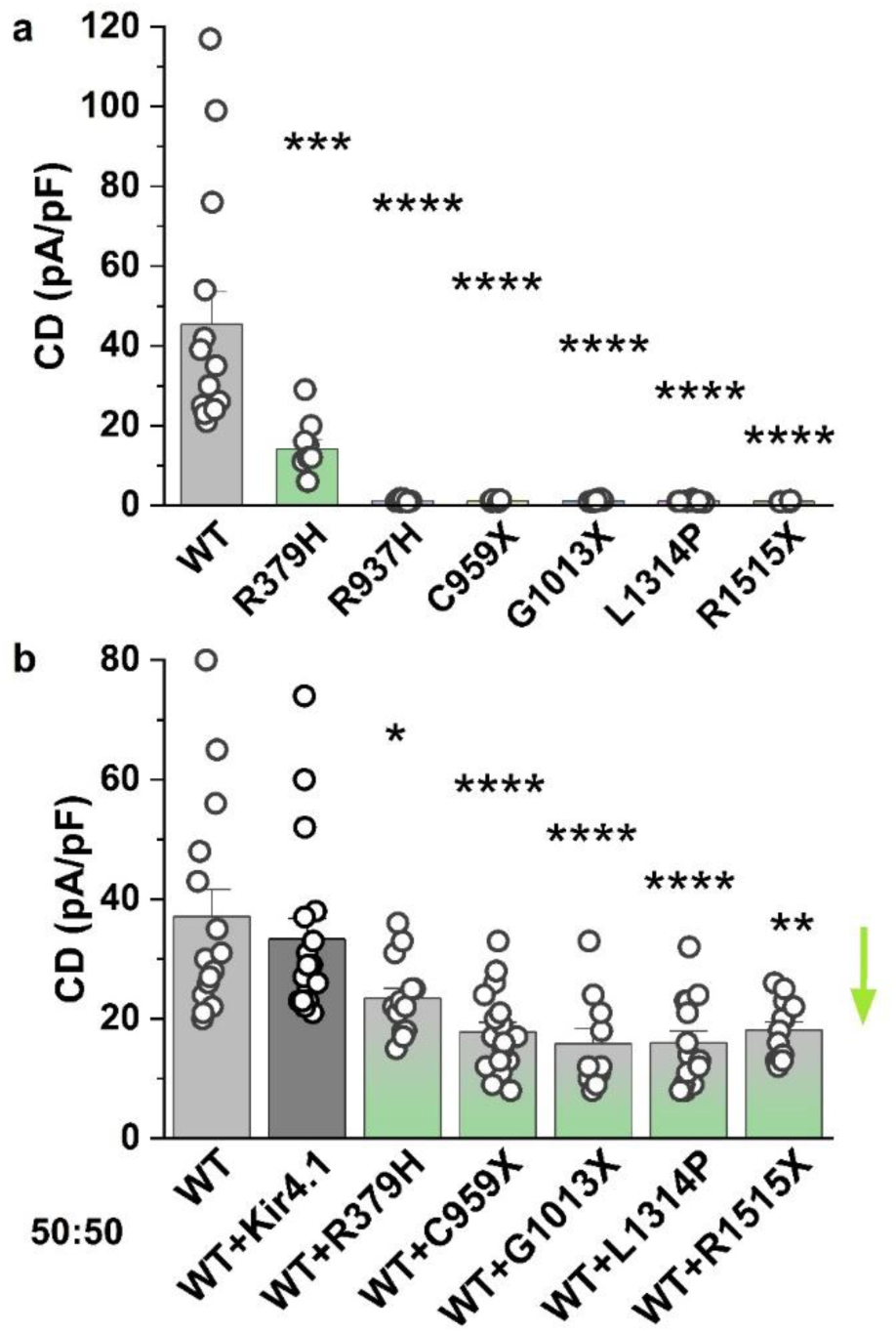
ASD hNav1.2 mutants are not rescued in neocortical neurons and show dominant negative effects. Maximal current density obtained from recordings of neocortical neurons in primary culture transfected with WT hNa_v_1.2 or the ASD mutant channels (**a**) or co-transfected with the WT and the mutant channels (**b**), which reduce the current density of the WT (the arrow highlights the negative dominant effects). The data on hNa_v_1.2-WT channels in control for panel b were obtained from a novel series of transfections performed in parallel with those of the other conditions displayed in the panel. Data shown as mean±SEM. See Supplementary material (Statistical Tables) for values, n and statistical tests.

### The negative dominant effect of the nsASD Na_v_1.2 mutants depends on regions that have been implicated in the interaction between Na^+^ channel α-subunits

To investigate the molecular basis of the dominant-negative effects produced by nsASD-associated *SCN2A* variants, we tested whether these could arise from physical interactions between Na_v_ α-subunits. Prior work, largely on the cardiac channel Na_v_1.5, suggested that Na_v_ α-subunits form functional dimers through binding of 14-3-3 to two sites within the DI–DII intracellular linker, and a second region in the same linker that mediates direct α-subunit interactions (residues 493–517in Na_v_1.5) independently of 14-3-3^22^ (Fig.4a). Inhibiting 14-3-3 action, either using the peptide blocker difopein or introducing the S460A mutation in Na_v_1.5 (which disrupts 14-3-3 binding), prevented dimer formation^22^.

**Figure 4:**
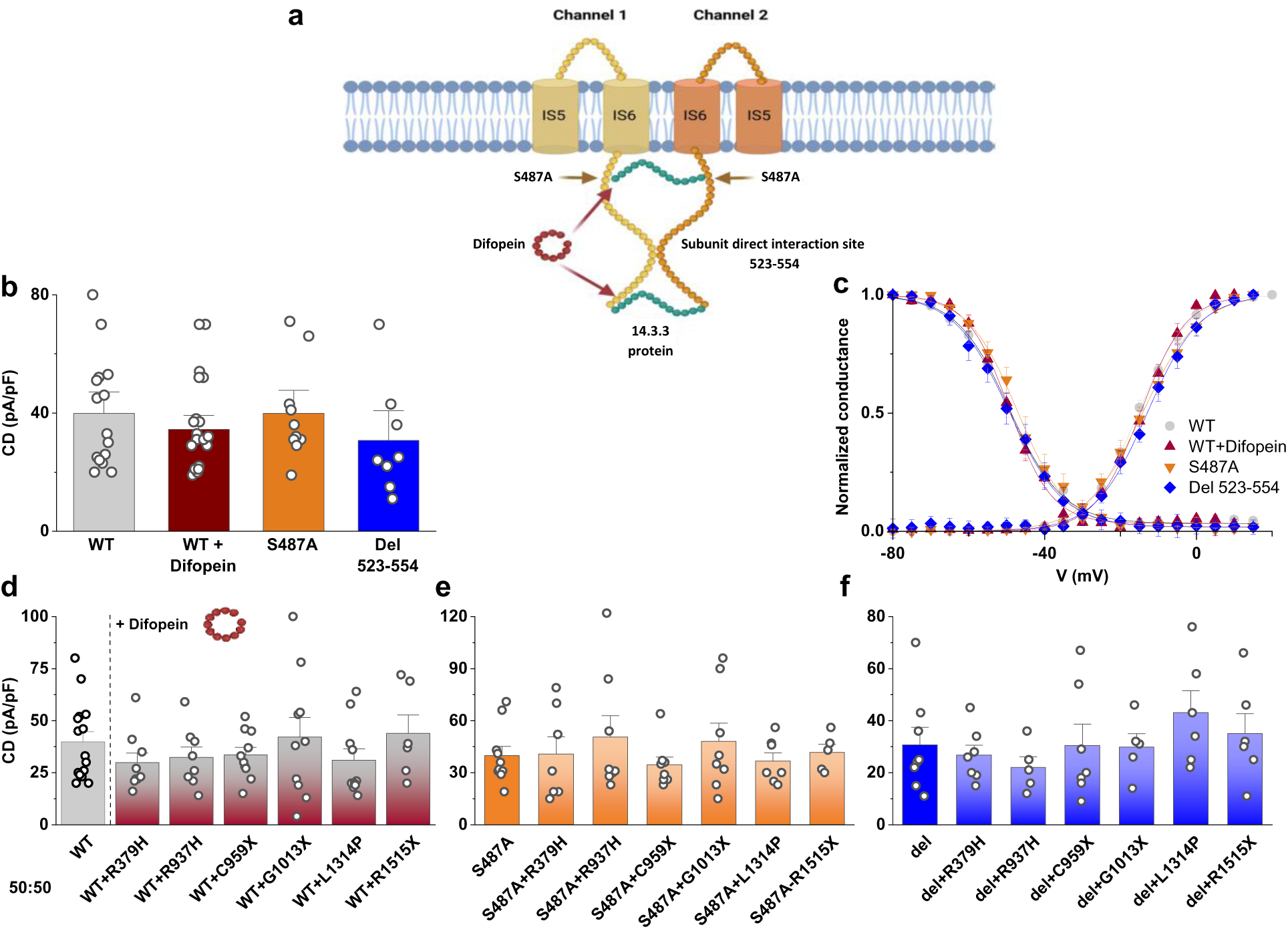
Domains that have been implicated in the interaction between two α subunits are necessary for the dominant negative effect of ASD Na_v_1.2 mutants. **(a)** Schematic illustration of the interaction between two sodium-channel α-subunits, in which the DI–DII intracellular linker engages 14-3-3 proteins to form an inter-subunit bridge while a defined interaction domain mediates direct contact and promotes subunit association (modified from reference ^22^). (**b**) Maximal current density of hNa_v_1.2-WT expressed in neocortical neurons in culture compared to that obtained in conditions that should inhibit the interaction (hNa_v_1.2-WT co-expressed with difopein, hNa_v_1.2-S487A and hNa_v_1.2-Del523-554). The data on hNa_v_1.2-WT channels in control were obtained from a novel series of transfections performed in parallel with those of the other conditions displayed in the figure. (**c**) Mean voltage dependence of activation and fast inactivation; lines are Boltzmann curves generated using the mean parameters of the fits of the single cells. (**d**) Maximal current density obtained in neocortical neurons co-transfected with hNa_v_1.2-WT and ASD mutant channels in the presence of difopein, compared with hNa_v_1.2-WT alone. (**e**) Maximal current density of neocortical neurons co-transfected with hNa_v_1.2-S487A and ASD mutant channels. (**f**) Maximal current density of neocortical neurons co-transfected with hNa_v_1.2-Del523-554 and ASD mutant channels. Data shown as mean±SEM. See Supplementary material (Statistical Tables) for values, n and statistical tests.

We tested whether the dominant-negative effects of nsASD-associated *SCN2A* variants require regions implicated in Na_v_ α-subunit dimerization by: (i) inhibiting 14-3-3 through difopein co-expression, (ii) introducing the S487A mutation in hNa_v_1.2 (analogous to S460A in Na_v_1.5), and (iii) generating the Δ523–554 deletion in hNav1.2, corresponding to the direct α–α interaction site identified in Na_v_1.5.

Control experiments evaluating the features of WT-hNav1.2 co-expressed with difopein, of hNav1.2-S487A, or of hNav1.2-Δ523–554 showed no changes in current density or in the voltage dependence of activation or fast inactivation compared to WT-hNav1.2 (Fig.4b–c), indicating that these manipulations did not alter main channel properties.

We then assessed their impact on the dominant-negative effects of nsASD mutants (Fig.3d–f). Inhibition of 14-3-3 with difopein abolished the reduction in current density normally produced by ASD mutants, and the S487A mutation produced the same rescue (Fig.4e). Thus, preventing 14-3-3 binding, which has been implicated in α-subunit dimerization, eliminates dominant-negative effects. Likewise, ASD mutants failed to reduce current density when co-expressed with Δ523–554, which lacks the site proposed to be implicated in direct α–α interaction (Fig.4f).

Together, these results support a model in which dimerization of Na_v_ α-subunits are required for nsASD-associated dominant-negative effects.

### The LoF and the negative dominant effects of nsASD mutants are caused by decreased plasma membrane targeting

It has been proposed that LoF of Nav1.2 nsASD mutants is caused by a functional impairment (non-conductive mutants) of channels that are efficiently targeted to the plasma membrane^13^. However, our data are consistent with a trafficking defect of WT-nsASD mutant α subunit dimers, with the reduction of plasma membrane targeting of the dimer due to increased retention in the endoplasmic reticulum and subsequent degradation, as also proposed for Na_v_1.5 mutants^23^.

To investigate if dominant negative effects observed evaluating the current density correlate with defects in cell surface expression, we performed binding experiments on intact transfected tsA201-cells expressing WT or ASD mutant channels using the radiolabeled α-scorpion toxin AaHII (^125^I-AaHII), which binds specifically with nanomolar affinity to the receptor site 3 of Na_v_ α subunits (the extracellular loop between S3-S4 transmembrane segments of domain IV)^36^. These binding experiments allow a sensitive and specific quantification of Na^+^ channel surface expression that is not contaminated by intracellular signals^34,35^. Our binding data show a tight correlation between the reduction of cell surface expression and the reduction of current density (Fig.5a). In fact, for the mutant R379H, both the current density and specific ^125^I-AaHII binding were reduced by 70-80%, whereas for R937H and L1314P, which do not generate current, no binding was observed. The truncating variants C959X, G1013X and R1515X do not retain the AaHII binding site and, as expected, show no signal in binding experiments (not shown). Thus, the reduction or absence of current that we observed for the ASD mutants in patch-clamp experiments are caused by a reduction or absence of plasma membrane targeting.

**Figure 5:**
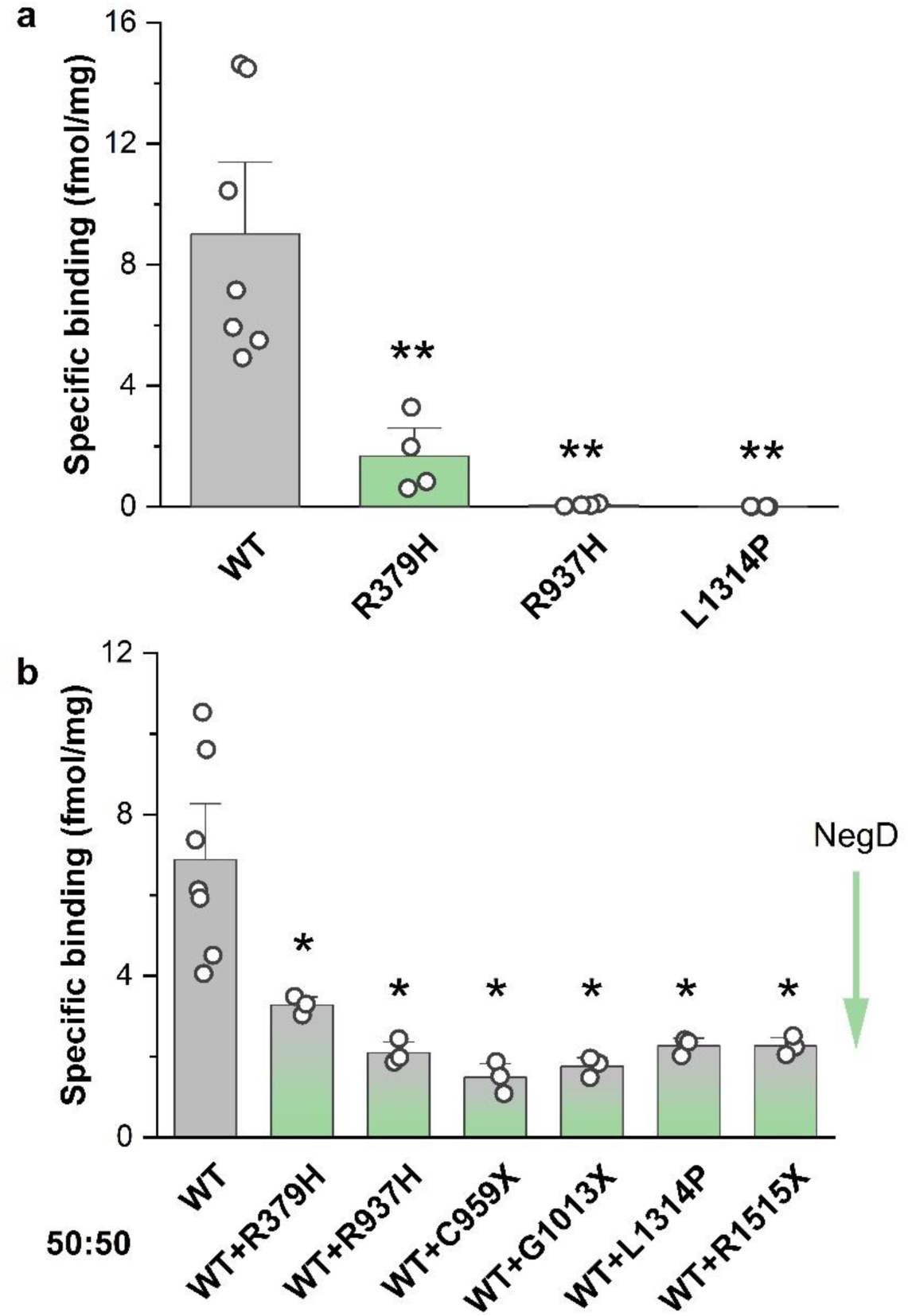
ASD mutants have decreased/lack of plasma membrane targeting and decrease membrane targeting of hNa_v_1.2-WT when co-expressed. **(a)** Specific binding of ^125^I-AaHII-scorpion toxin to intact tsA-201 cells transfected with hNa_v_1.2-WT or the mutants that retain the binding site of the toxin. **(b)** Specific binding of ^125^I-AaHII-scorpion toxin to intact tsA-201 cells co-transfected with hNa_v_1.2-WT and the ASD mutants, compared with cells transfected with hNa_v_1.2-WT alone (50% cDNA in the transfection); in this assay, binding reflects only hNav1.2-WT channels when co-transfected mutants lack the AaHII binding site. The arrow highlights the negative dominant effect. Data shown as mean±SEM. See Supplementary material (Statistical Tables) for values, n and statistical tests.

Next, we performed binding experiments on intact tsA-201 cells co-transfected with WT and nsASD mutants to reproduce heterozygous conditions. In this assay, binding reflects only hNav1.2-WT channels when co-transfected mutants lack the AaHII binding site. Our data show that the channels targeted at the cell surface was reduced by >50% when the nsASD mutants are co-expressed (Fig.5b). Therefore, our binding data correlate with the results of patch-clamp experiments and demonstrate that the targeting to the membrane of hNa_v_1.2-WT is drastically reduced when nsASD mutants are co-expressed. Overall, our patch-clamp and binding data show that the ASD mutants induce dominant negative effects by reducing the targeting to the cell surface of WT-channels, which is mediated by the interaction between WT and nsASD mutant α subunits.

### hNa_v_1.2 mutants causing other phenotypes with infantile-childhood onset do not show dominant negative effects

We compared the mechanism of action of ASD mutants with that of other LoF hNa_v_1.2 mutants. We studied hNa_v_1.2 mutations causing other phenotypes with clinical onset in the infantile-childhood period or later, such as ICDEE (D284G, G659D, A896V, C1344Y and R1882X), schizophrenia (R850P and V1289F), and ASD with severe epileptiform EEG activity without clinical seizures (T1420M and R1635Q; although for T1420M we inferred it considering the functional study). Fig.6a shows representative Na^+^ current traces elicited with a series of depolarizing steps in tsA-201 cells transfected with WT or mutant channels. The analysis of the maximal current density is shown in Fig.6b and indicates that all the mutants studied (except D284G) induced a significant reduction of current, although none of them show complete LoF. To find out if the cellular background could influence the expression at the cell surface, we transfected them in neocortical neurons and performed patch-clamp experiments to measure the maximal current density, as we did for ASD mutants (Fig.6c), observing that R1635Q was rescued by the neuronal cellular background. Therefore, differently from ASD mutants, all these mutants can be targeted to the plasma membrane and can generate currents; some of them are rescued by the neuronal cellular background.

**Figure 6:**
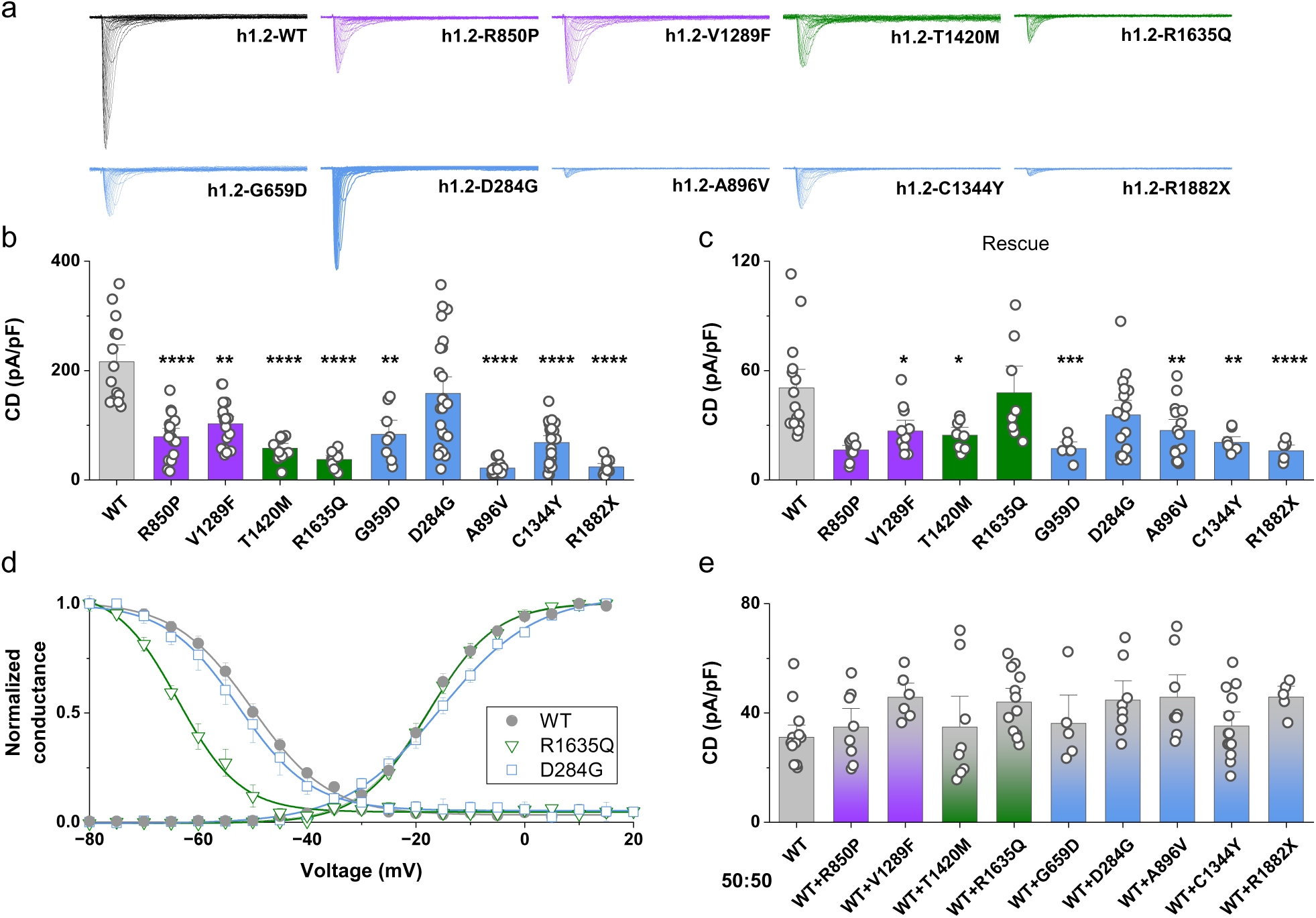
Functional analysis of hNa_v_1.2 mutants responsible for other phenotypes with infantile-childhood onset. **(a)** Representative whole-cell Na^+^ current families for hNa_v_1.2 WT and hNa_v_1.2 mutants responsible for schizophrenia (R850P and V1289F), ASD with epileptiform EEG activity without clinical seizures (T1420M and R1635Q) and DEE (D284G, G659D, A896V, C1344Y and R1882X) expressed in tsA-201 cells. Maximal current density of the WT and the different mutants expressed in tsA-201 cells **(b)** or expressed in neocortical neurons in culture (**c**). (**d**) Voltage dependence of the hNa_v_1.2-WT and mutants. (**e**) Maximal current density of the hNa_v_1.2-WT co-expressed with the different mutants in neocortical neurons. Data shown as mean±SEM. See Supplementary material (Statistical Tables) for values, n and statistical tests.

To better characterize the functional impact of R1635Q, which is rescued in neurons, as well as of D284G, which does not show a reduction of current density in either tsA-201 cells or in neurons, we studied their voltage dependence of activation and inactivation in transfected neocortical neurons (Fig.6d). The R1635Q variant induced a large 10.8-mV negative shift of the voltage dependence of inactivation, indicating a clear LoF. The voltage dependence of activation was less steep for D284G possibly indicating a mild LoF, although the half maximal potential was not modified.

We also studied the effect induced by the co-expression in neurons of hNa_v_1.2-WT with the mutants involved in DEE, schizophrenia and ASD with epileptiform EEG activities without clinical seizures. None of the mutants modified the current density of the co-expressed WT channel (Fig.6e), not even those exhibiting a dramatic reduction in current density when expressed alone (R850P, T1420M, G659D, C1344Y and R1992X).

Although the analysis of current density and voltage dependencies was consistent with LoF for all mutants, these parameters represent only a subset of functional properties. A comprehensive assessment of gating properties is labor-intensive, and integrating multiple functional alterations into a single, interpretable functional outcome is often not straightforward. To disclose the overall impact of these variants, we therefore recorded in transfected neurons action Na^+^ currents elicited in response to a neuronal discharge, as performed in Fig.2c. However, because the action potential-evoked Na^+^ currents were already small in WT hNav1.2 under our recording conditions, potential LoF effects of some of the variants could not be reliably resolved. To overcome this limitation, we enhanced the current by adding the sea anemone toxin ATX–II (10 nM) to the extracellular solution. ATX–II slows fast inactivation of voltage–gated Na^+^ channels, thereby increasing the current during repetitive action potential firing, while exerting only small effects on the kinetics of activation. Representative traces of hNa_v_1.2-WT with or without 10 nM ATX II are shown in Fig.7a, and action currents obtained in response to an action potential discharge, displayed as mean current densities, are shown in Fig.7b. Using these conditions, we studied the overall functional effect of the variants implicated in schizophrenia, ASD with epileptiform EEG activity and DEE, as well as of the only nsASD mutant that showed residual current when expressed alone (R379H). The analysis of the peak current density of the first action current is shown in Fig.7c and indicates that its amplitude is significantly reduced for all the mutants tested, except for R1635Q and D284G. However, comparing the mean peak amplitude of the last three action currents in the discharge, we found that all mutants induced a significant decrease in comparison to WT (Fig.7d). These data point to LoF as the general effect of all the mutants studied, which is consistent with neuronal hypoexcitability.

**Figure 7:**
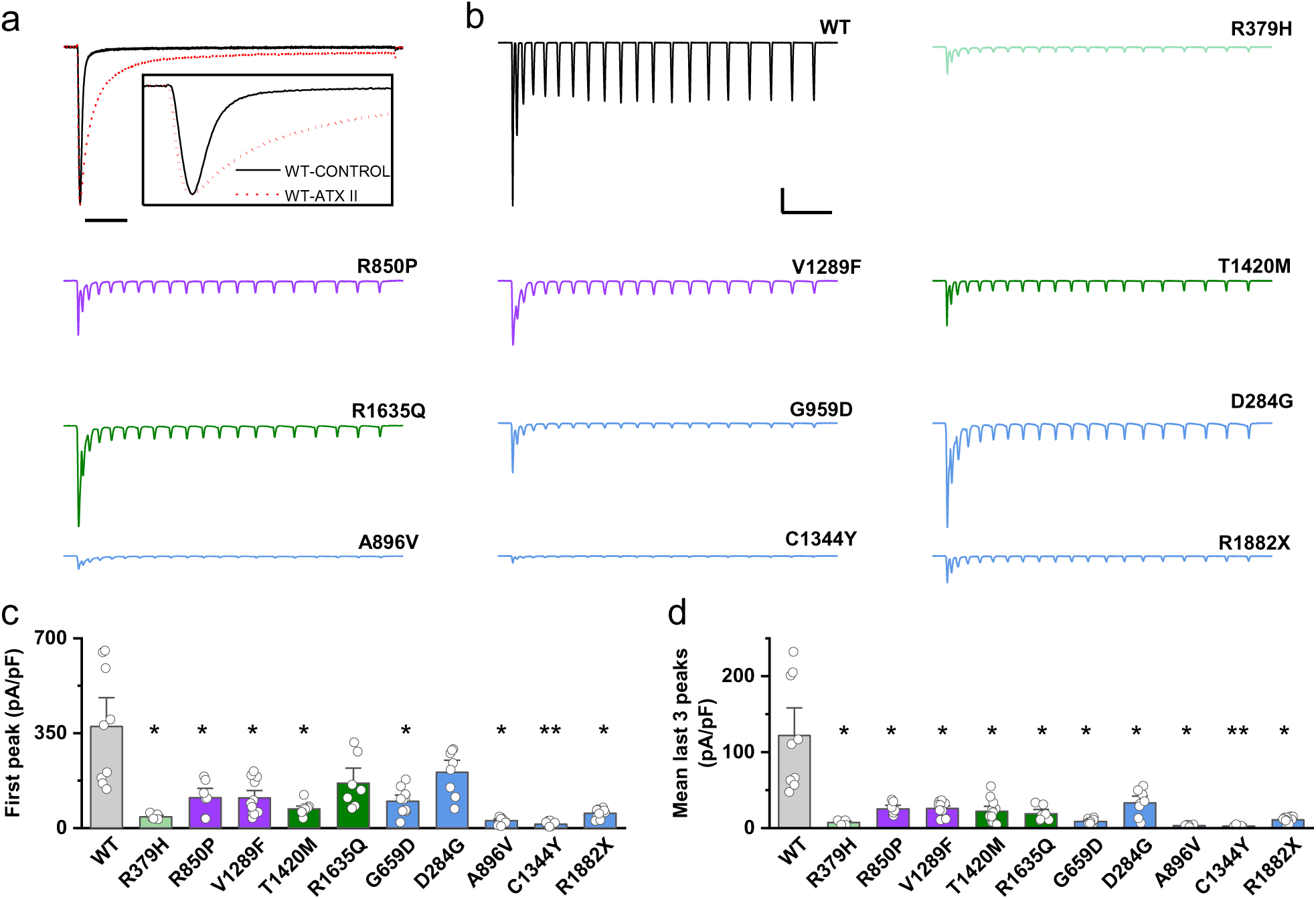
Overall effects of hNav1.2 variants evaluated with action potential discharge-clamp in the presence of ATX II. **(a)** Representative normalized hNa_v_1.2 Na^+^ current traces elicited by a step depolarization at the maximal peak current density in control or in the presence of 10 nM ATX II; scale bar 20ms. The inset shows the same traces in a 7-ms window. **(b)** Action Na^+^ currents in the presence of ATX II displayed as mean of current densities for hNa_v_1.2-WT, hNa_v_1.2-R379H (nsASD with residual current), hNa_v_1.2-R850P and hNa_v_1.2-V1289F (schizophrenia), hNa_v_1.2-T1420M and hNa_v_1.2-R1635Q (ASD with epileptiform EEG discharges), hNa_v_1.2-G659D, hNa_v_1.2-D284G, hNa_v_1.2-A896V, hNa_v_1.2-C1344Y and hNa_v_1.2-R1882X (infantile-childhood DEE); scale bars 50pA/pF, 25ms. Comparison of the current density of the first action current **(c)** and of the mean of the last three action currents in the discharge **(d)**. Data are shown as mean ± SEM. See Supplementary material (Statistical Tables)for values, n and statistical tests.

Only nsASD variants exhibited negative dominant effects, a pattern that was statistically significant (Fisher’s exact test, p = 0.0002). This indicates that the mechanism of nsASD variants observed in heterologous systems is distinct from that of other LoF variants.

## DISCUSSION

### *All the SCN2A* variants tested induce a loss of function of hNa_v_1.2 channels in both cell lines and cultured neurons

All *SCN2A* variants characterized in our study, whether identified from the literature or newly reported, induced Na_v_1.2 LoF. This shared functional effect was independent of the associated clinical phenotype, which was within the late infantile or childhood onset spectrum for all. The LoF resulted from reduced peak Na^+^ currents or altered biophysical properties, as observed for D284G, A896V, and R1635Q. Variants associated with nsASD exhibited complete LoF, with the exception of R379H. The R379H variant recapitulates the effect of the homologous R367H variant in Na_v_1.5, which causes Brugada syndrome and shows a large reduction in peak current density^42^.

Our functional data show prominent LoF for nsASD variants, consistent with other studies^13^. Two hypotheses have been proposed to explain the absence of measurable current in these variants: (1) the mutant channels reach the plasma membrane but fail to conduct, or (2) trafficking to the plasma membrane is impaired. Immunohistochemistry and TIRF microscopy have been used to propose that ASD mutants localize to the plasma membrane but are non-conducting^13^. However, these measurements can be contaminated by signal from the endoplasmic reticulum. To further address this question, we used a different approach: binding assays with a radio-labelled scorpion toxin on intact transfected cells, which is a highly quantitative, sensitive, and specific measure of the density of functional channels at the plasma membrane, offering a more direct readout of specific surface expression than other methods. Our results showed no detectable binding for the nsASD mutants tested, but for R379H that showed reduced binding, consistent with the reduction in current density observed in patch-clamp recordings. These findings suggest that nsASD mutants fail to traffic properly to the plasma membrane, rather than inducing impaired conductance.

### Heterozygous conditions highlight different mechanisms of action between nsASD *SCN2A* variants and *SCN2A* variants underlying other phenotypes

To better model the heterozygous pathophysiological state, we co-expressed WT and mutant Na_v_1.2 channels. The nsASD-associated mutants consistently diminished WT channel function by approximately 50%, as evidenced by reductions in both Na^+^ current density, measured via patch-clamp recordings, and surface expression, assessed through binding assays. These findings indicate a dominant-negative mechanism. By contrast, variants linked to other phenotypes did not show such dominant-negative effects. This finding aligns with the more pronounced autistic phenotype reported in mouse models exhibiting greater reductions in *Scn2a* expression^18–21^, relative to haploinsufficient *Scn2a*^+/-^ mice^3,14–17^.

The R1635Q variant identified in our study was initially classified as nsASD, but it does not exert a dominant-negative effect. However, upon completion of the clinical evaluation, this variant was found to be associated with a more complex phenotype, including global cognitive regression, absent speech, and continuous centrotemporal spikes during sleep, without clinical seizures, indicating a presentation inconsistent with nsASD. Similarly, the T1420M variant, previously reported in association with nsASD^39^, shows functional properties comparable to R1635Q. Given the lack of detailed phenotypic data for the T1420M carrier, we propose reclassifying this variant alongside R1635Q. The patient carrying the dominant-negative L1314P variant, who underwent the same comprehensive clinical assessment, did not exhibit significant EEG abnormalities. Collectively, these findings suggest that a dominant-negative effect correlates with pure nsASD phenotypes, while its absence is associated with different clinical presentations.

### Domains implicated in the dimerization of hNa_v_1.2 channels are necessary for the dominant negative effect of ASD mutations

Previous studies have proposed that dominant-negative effects of the cardiac Na^+^ channel Na_v_1.5 can arise from interactions between wild-type (WT) and mutant Na^+^ channel α-subunits^22,23,43^. The proposed interaction site lies within the IS6–IIS1 linker, which contains both a 14-3-3 protein binding motif and a region between amino acids 523–554 mediating a direct α-subunit interaction.

We sought to directly test the dimerization hypothesis as a mechanism underlying dominant-negative effects of Na_v_1.2 mutants. We employed several complementary strategies: inhibition of 14-3-3 binding using difopein, mutation of serine 487 to alanine (S487A) in hNa_v_1.2 to block 14-3-3 association, and deletion of the region proposed to mediate direct α subunit interactions (Δ523–554). Each of these conditions abolished the dominant-negative effect in our experiments, providing strong evidence that this phenomenon depends on the previously proposed α-subunit interaction site. Moreover, co-transfection of WT channels with nsASD-associated variants revealed that these mutants reduce trafficking of the WT-mutant complex to the plasma membrane, as quantified by radioligand binding assays. Together, these findings indicate that nsASD-associated mutations exert their effects through a dominant-negative mechanism that requires the proteins and domains that have been implicated in interaction between α-subunits. Our binding studies further reveal that, unlike correctly folded channels that are trafficked to the plasma membrane, the reduced targeting of WT-nsASD or nsASD-nsASD complexes to the plasma membrane results in the intracellular retention of the dimer, which is probably subsequently degraded (Fig.8).

**Figure 8:**
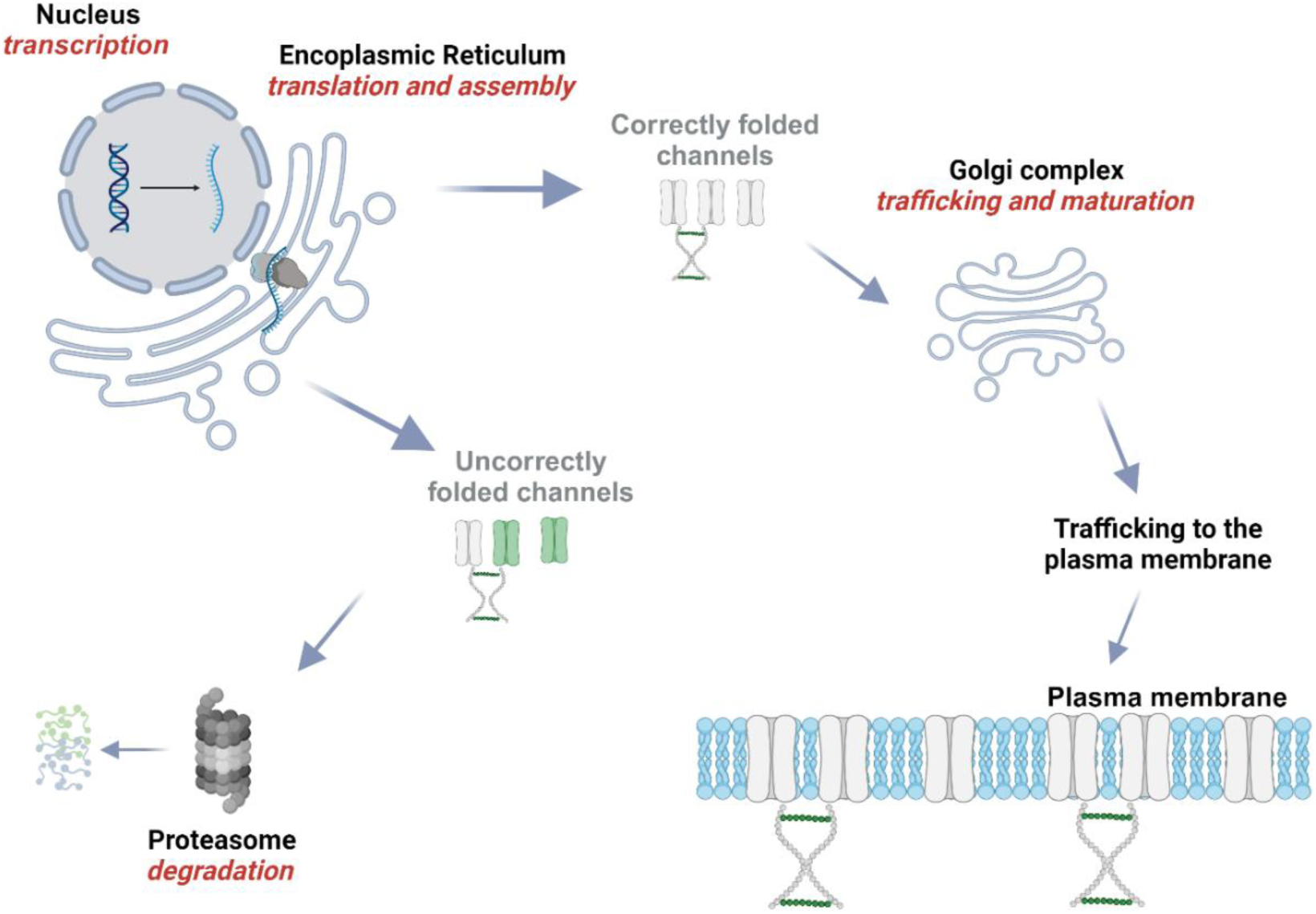
Proposed mechanism of action of non-syndromic ASD (nsASD) hNa_v_1.2 mutants. We propose that nsASD-linked mutations act through a dominant-negative mechanism that depends on interactions between α-subunits. WT–nsASD and nsASD–nsASD dimers exhibit reduced surface trafficking: they are retained intracellularly, where they are likely targeted for degradation.

Despite reduced current density, dominant-negative effects were not observed for variants associated with DEE, schizophrenia and ASD with continuous epileptiform EEG discharges during sleep. Overall, our data reveal a specific mode of action for ASD-associated *SCN2A* variants and highlight potential avenues for targeted therapeutic interventions.

Our findings for *SCN2A* variants are part of a broader pattern observed in voltage-gated ion channels, in which certain truncating or missense mutations can exert dominant-negative effects on wild-type subunits, involving aberrant interactions between truncated or misfolded subunits and wild-type channels that lead to retention in the endoplasmic reticulum and subsequent degradation. This mechanism has been proposed for the homologous cardiac Na_v_1.5 channel, where Brugada syndrome-associated truncations reduce wild-type channel function both in vitro and in vivo^44^. Similarly, truncating mutations in Cav2.1 (P/Q-type) channels associated with episodic ataxia type 2 (EA2) can suppress wild-type channel currents through similar mechanisms of ER retention/impaired trafficking, and *in vivo* studies in EA2 knock-in mice support the pathological relevance of these interactions, consistent with a dominant negative effect^45,46^.

Some of the *SCN2A* mutants showing dominant negative effects are truncated proteins. Although premature termination codons located far upstream of exon–exon junctions are generally expected to trigger nonsense-mediated mRNA decay (NMD), incomplete mRNA degradation can allow synthesis of truncated mutants^47,48^, which may exert dominant-negative effects by interacting with wild-type proteins in the endoplasmic reticulum. Such effects have been observed both *in vitro* and *in vivo* for Na_v_1.5 and Ca_v_2.1 truncations, where misfolding, ER retention, and proteasomal degradation can reduce wild-type channel function^23,43,44,49^. These findings suggest that even small amounts of truncated channel subunits can interfere with trafficking or function of wild-type proteins, providing a mechanism for dominant-negative pathogenicity across Na^+^ and Ca^2+^ channelopathies. Notably, while dominant-negative effects are robustly observed in heterologous expression systems for multiple channel families, *in vivo* evidence is more nuanced. These observations highlight that dominant-negative effects can be context-dependent and may be modulated by variant position, NMD efficiency, ER quality control features, and expression levels^47,48^. Overall, the mechanism we identified for nsASD variants is consistent with the broader paradigm of dominant-negative effects observed across Na^+^ and Ca^2+^ channelopathies, reinforcing the importance of evaluating each variant in pathophysiological relevant systems to understand its pathogenic potential.

### Clinical relevance of the functional study

From a clinical perspective, functional data have potential direct implications for stratification and management of patients. Determining whether a variant acts through haploinsufficiency or a dominant-negative mechanism can be important not only for mechanistic interpretation but also for precision medicine.

Our findings support an association between molecular mechanisms and clinical phenotypes. We observed that patients carrying dominant-negative *SCN2A* variants present with ASD but no seizures or severe epileptiform activity, whereas variants causing haploinsufficiency or non-dominant LoF are more frequently associated with epilepsy, often configuring a DEE, or other phenotypes, although there was no clear correlation between amount of LoF and phenotype for these variants. Our data underscore how functional stratification of *SCN2A* variants could refine clinical classification and prognosis, helping to distinguish between pure neurodevelopmental disorders and epilepsy syndromes within the same genetic continuum.

A recent study demonstrated that upregulating the remaining functional *SCN2A* allele using CRISPR activation (CRISPRa) can rescue neurological phenotypes in *Scn2a* haploinsufficient mice, highlighting a potential therapeutic strategy for neurodevelopmental disorders associated with *SCN2A* haploinsufficiency^50^. However, in the presence of dominant-negative variants, the concomitant increased expression of mutant channels could counteract the benefits of such approaches. Consequently, future therapeutic strategies that enhance Na^+^ channel expression should integrate detailed clinical phenotyping with functional data, to guide variant-specific interventions and avoid potential adverse effects in dominant-negative conditions.

The translational relevance of our findings depends on validating the functional consequences of *SCN2A* variants in patients. Although our experiments in heterologous cells and rodent cortical neurons in culture reveal clear mechanistic distinctions between haploinsufficient and dominant-negative variants, the extent to which these mechanisms operate in human neurons remains to be fully defined. While patient-derived induced pluripotent stem cell (iPSC)–based neurons and cortical organoids may offer complementary insight, variability in differentiation state, cellular maturity, and line-to-line heterogeneity can limit the reproducibility and interpretability of results obtained from these systems^51^, which should be considered as informative but not definitive.

*In vivo* studies in animal models are warranted for establishing the relevance of dominant-negative effects and determining how *SCN2A* nsASD variants specifically influence neuronal circuits and behaviour within an intact organism during different stages of development.

Although additional studies are needed to better elucidate detailed molecular mechanisms underlying dominant-negative effects, and clarify why such effects are not observed in variants not associated with nsASD, our findings indicate that assessing negative dominant effects in heterologous systems may serve as a useful biomarker for predicting the clinical phenotype of LoF *SCN2A* variants.

## Conclusions

We analyzed *SCN2A* variants associated with infantile and childhood phenotypes and confirmed that all produce Na_v_1.2 loss of function. The nsASD–linked variants caused the most severe deficits, largely due to impaired trafficking, and uniquely showed dominant–negative effects when co–expressed with wild–type channels, consistent with interactions between mutant and WT α–subunits. This mechanism distinguishes ASD–related variants and may offer an *in vitro* biomarker, which should be validated in larger cohorts. Our findings underscore the importance of integrating functional data into diagnostic workflows for precision medicine in neurodevelopmental channelopathies.

## SUPPLEMENTARY MATERIAL

**Supplementary Table 1.**
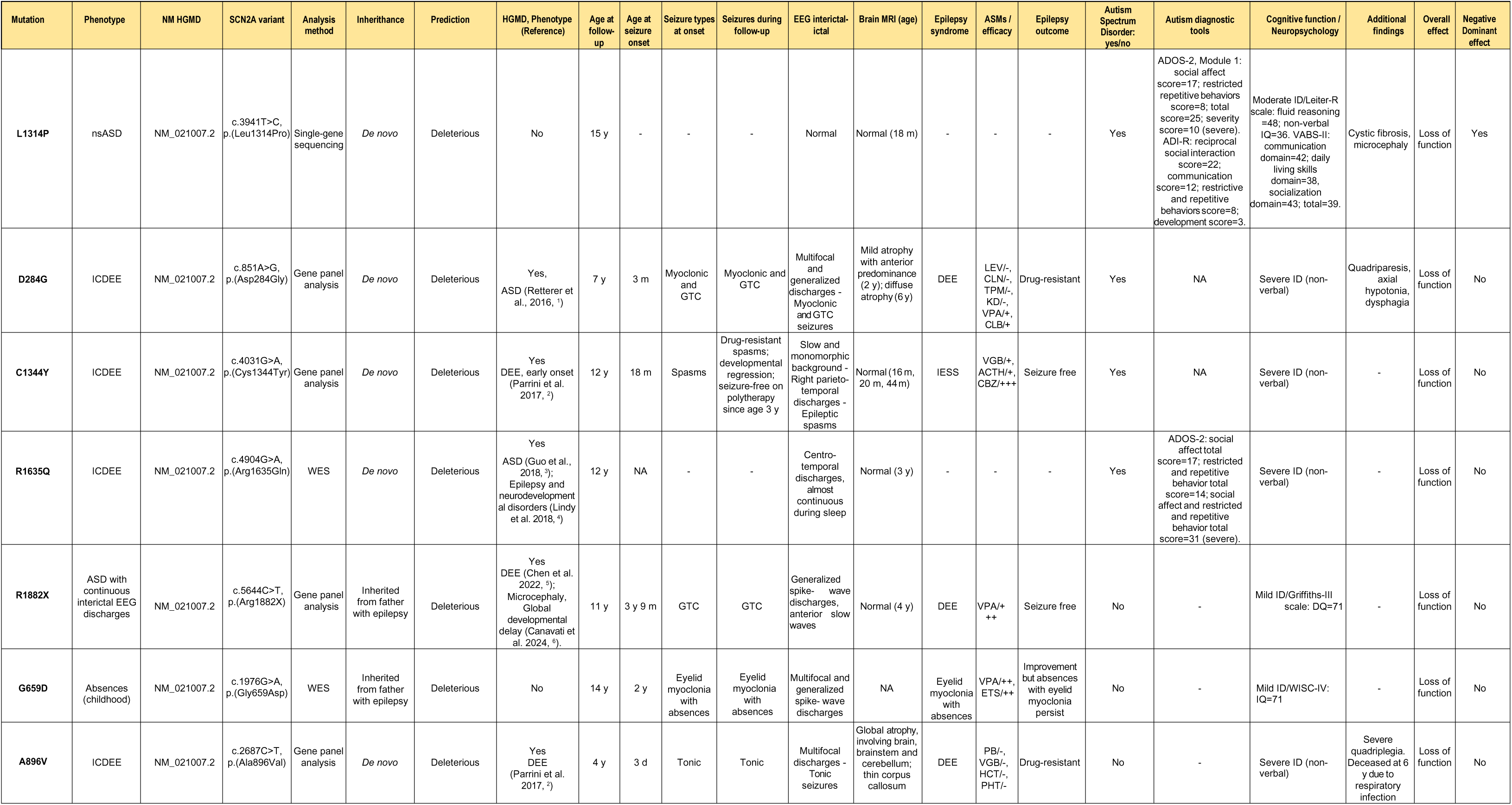
Clinical features of the patients from the cohort of the Meyer Children’s Hospital. Abbreviations: ACTH, adrenocorticotropic hormone; ADI-R, autism diagnostic interview–revised; ADOS, autism diagnostic observation schedule; ASD, autism spectrum disorder; ASMs, anti-seizure medications; CBZ, carbamazepine; CLB, clobazam; CLN, clonazepam; d, days; DEE, epileptic and developmental encephalopathy; DQ, developmental quotient; EEG, electroencephalogram; ETS, ethosuximide; GAI, general ability index; GTC, generalized tonic-clonic; ICDEE, infantile-childhood epileptic and developmental encephalopathy; HCT, hydrocortisone; HGMD, human gene mutation database; KD, ketogenic diet; ID, intellectual disability; IESS, infantile epileptic spasms syndrome; IQ, intelligence quotient; LEV, levetiracetam; m, months; NA, not available; nsASD, non-syndromic autism spectrum disorder; PB, phenobarbital; PHT, phenytoin; TPM, topiramate; VABS-II, vineland adaptive behavior scales second edition; VGB, vigabatrin; VPA, valproic acid; WES, Whole Exome Sequencing; WISC-IV, Wechsler intelligence scale for children-fourth edition; y, years; +, mildly effective; ++, effective; +++, very effective; -, ineffective.

---

**Supplementary Figure 1.**
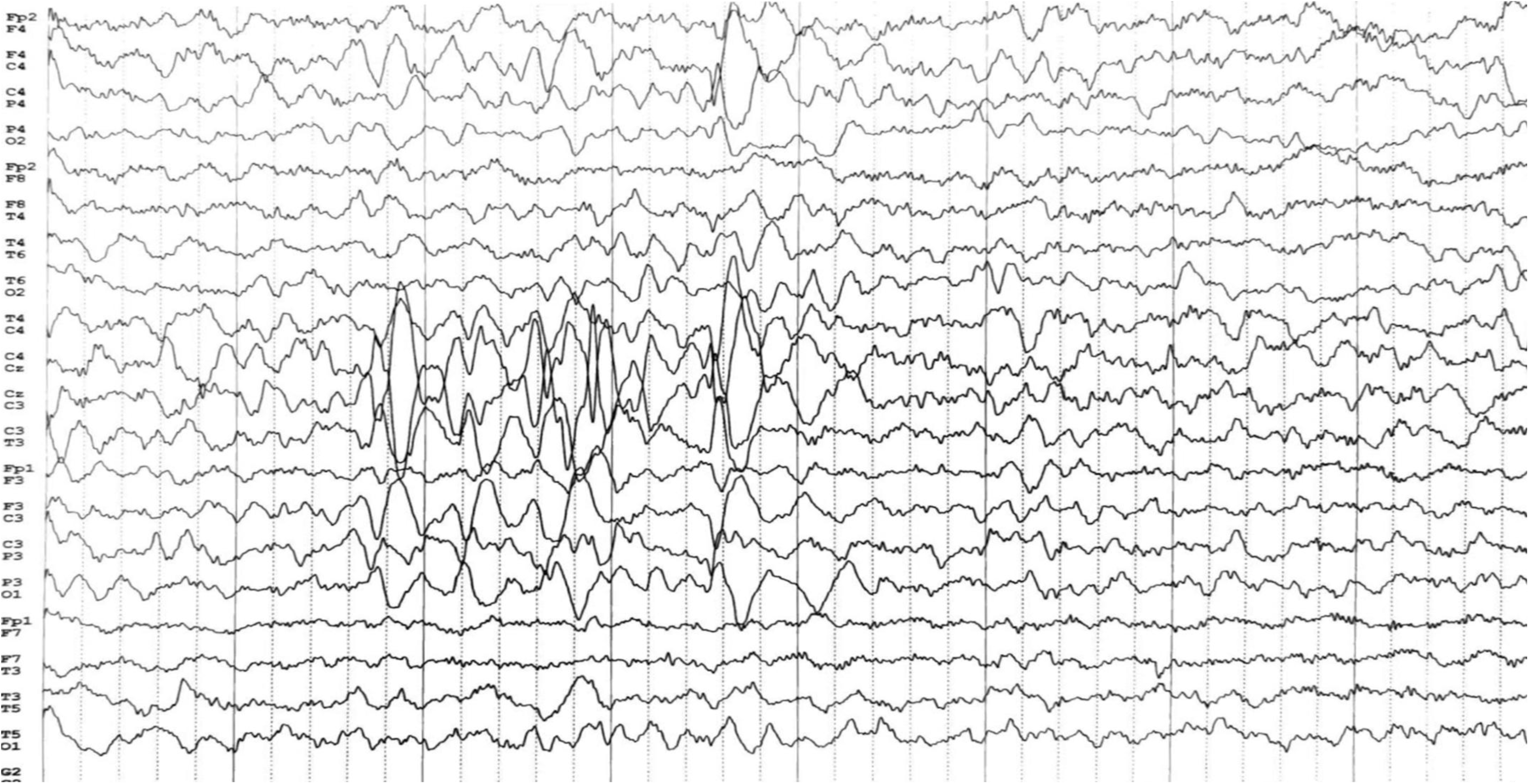
Patient carrying the L1314P variant. EEG recording while falling asleep shows sharp wave discharges over the vertex and central regions that are more pronounced than the physiological expected vertex sharp waves accompanying wakefulness-sleep transition.

**Supplementary Figure 2.**
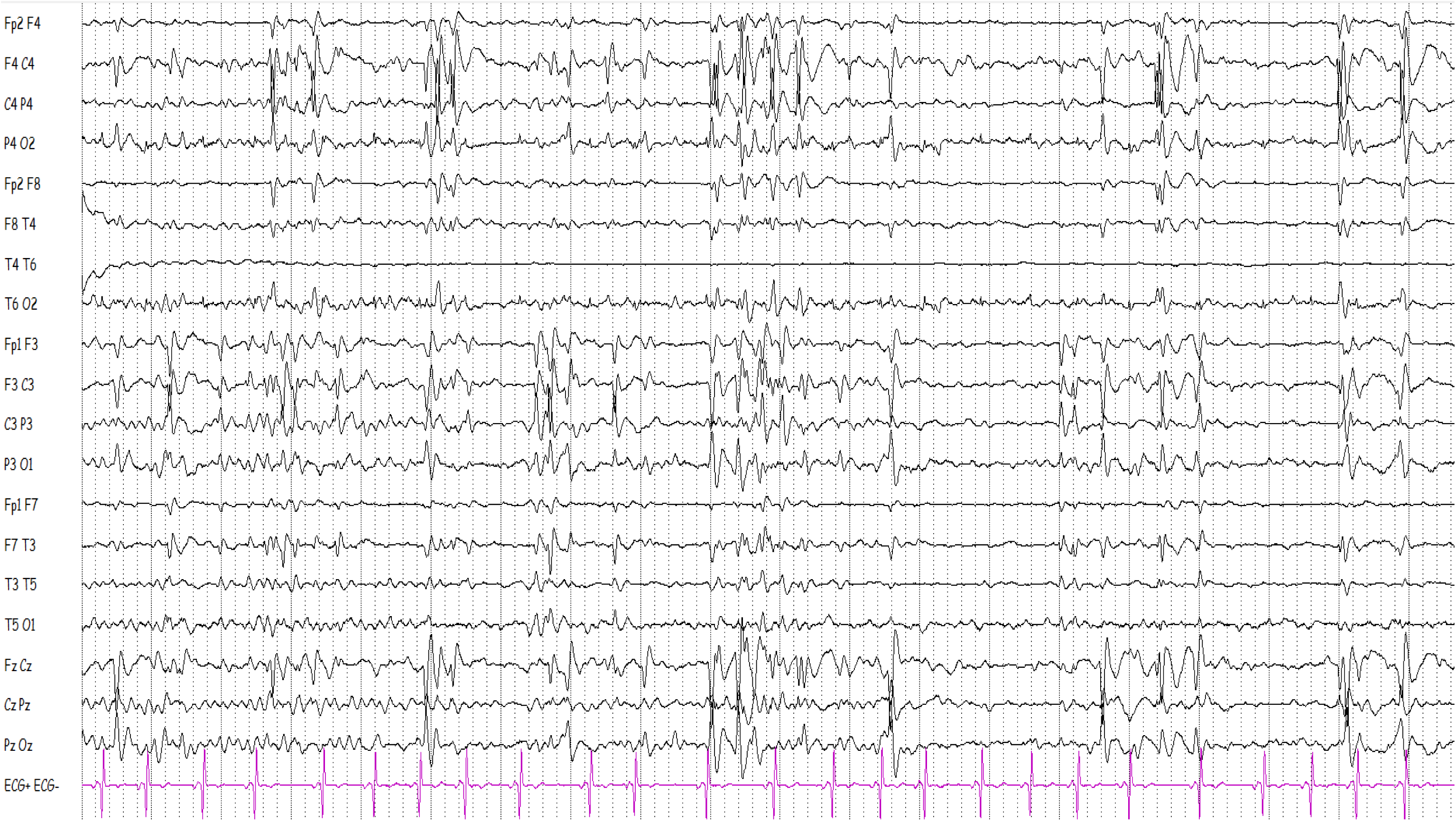
EEG recording during sleep of the patient carrying the R1635Q variant. There is activation of very frequent bilaterally synchronous and asynchronous spikes and sharp waves over the central regions and the vertex area.

## Statistical tables: values, n and statistical tests for main figures

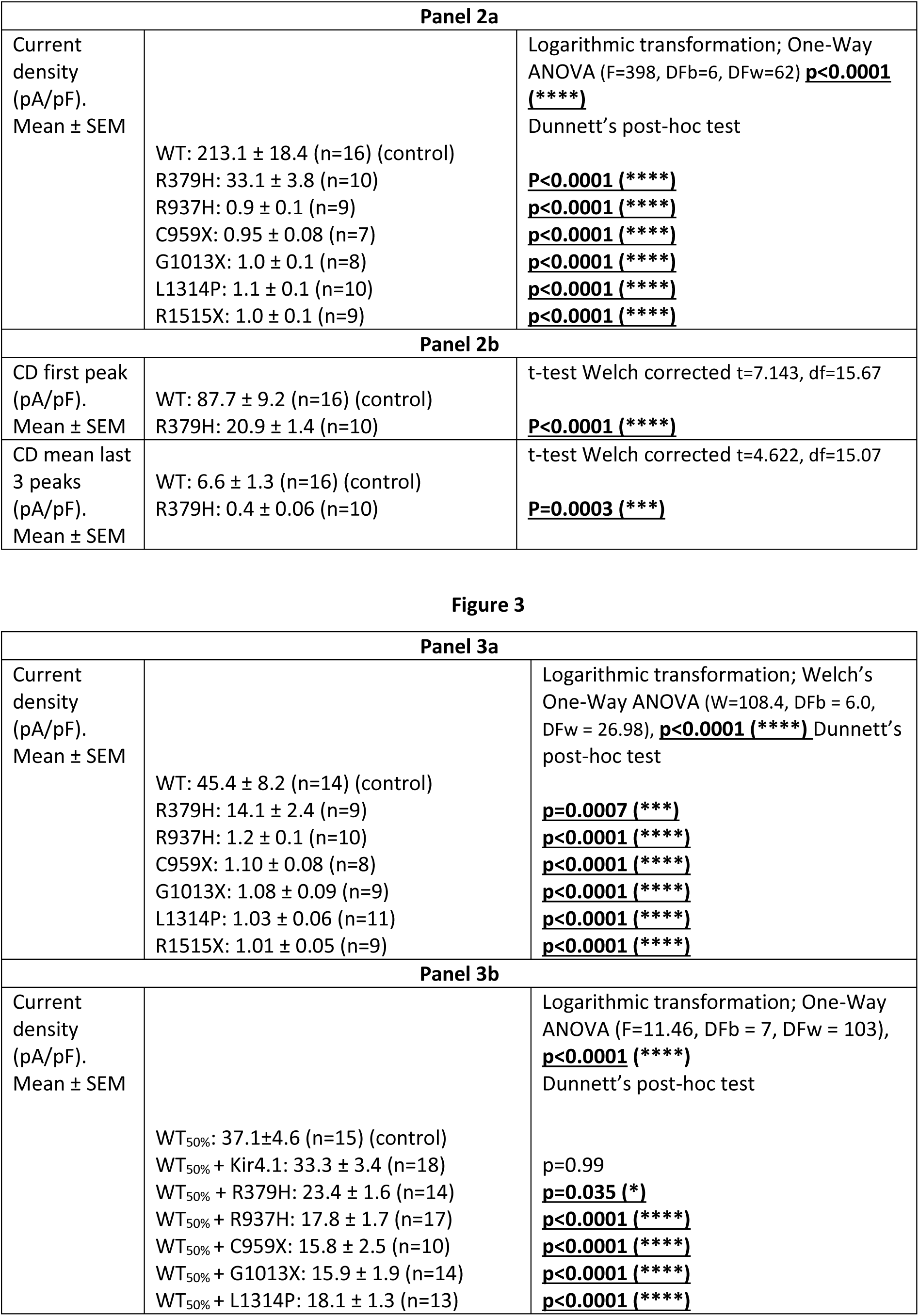
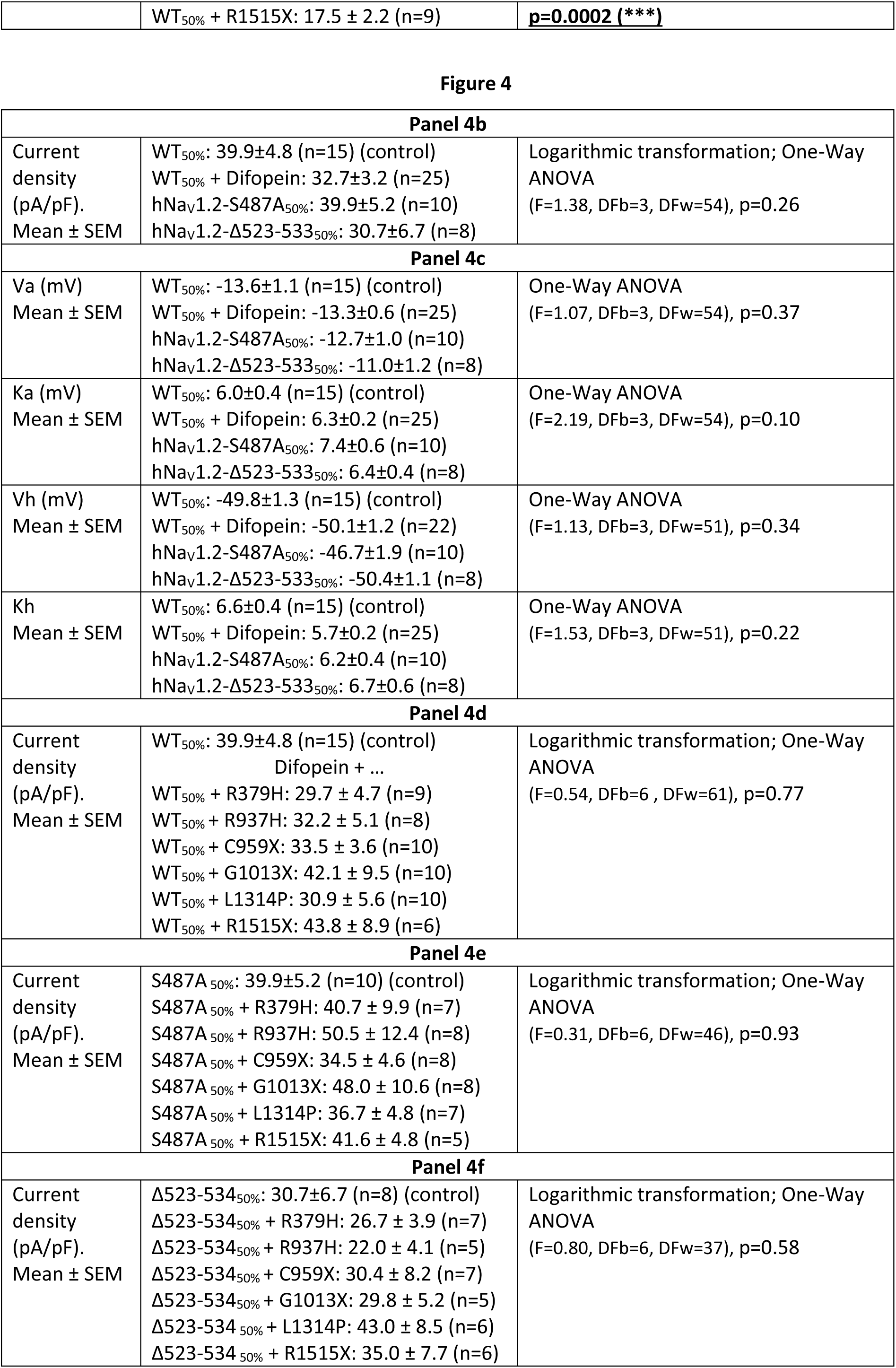
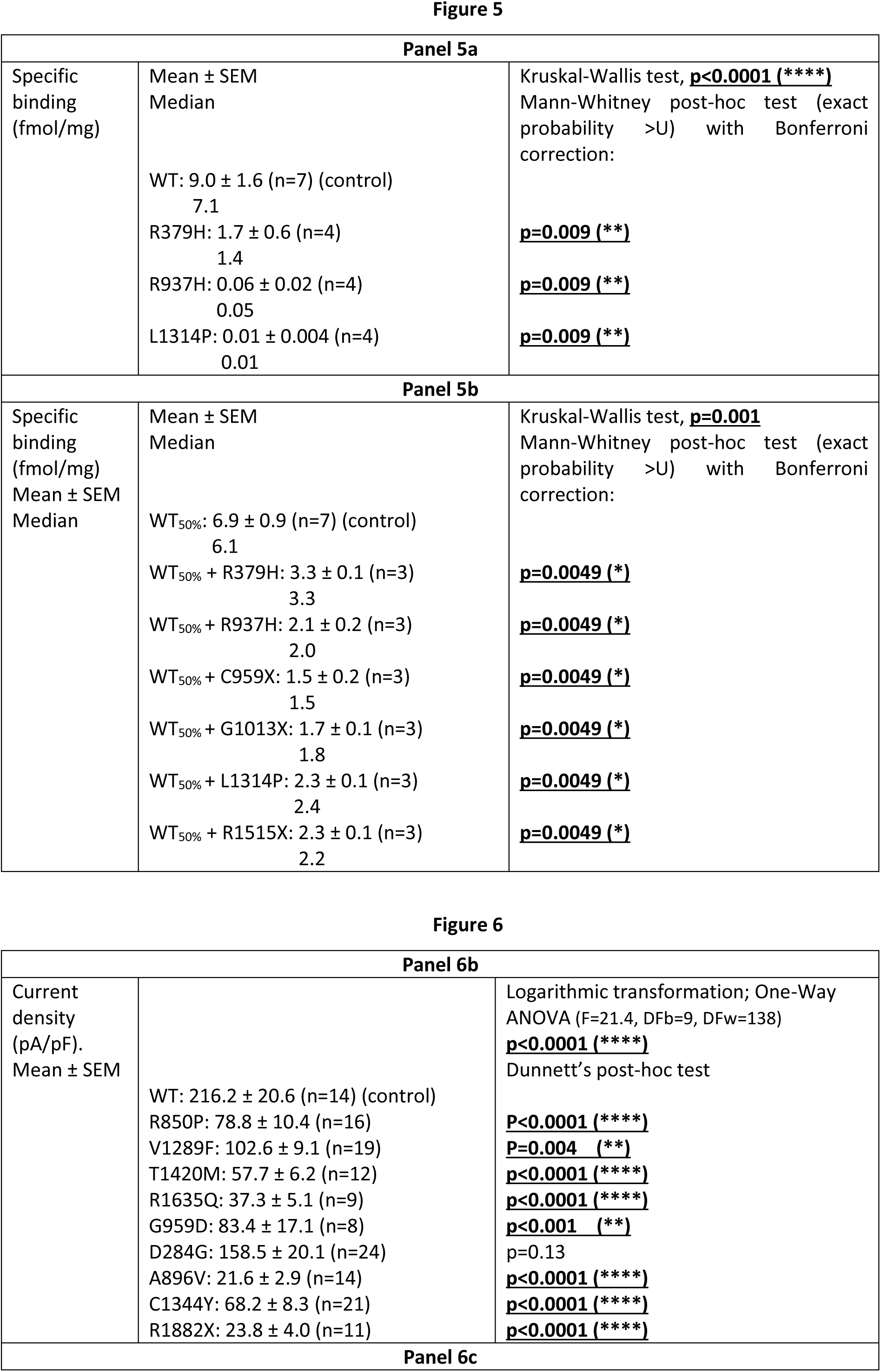
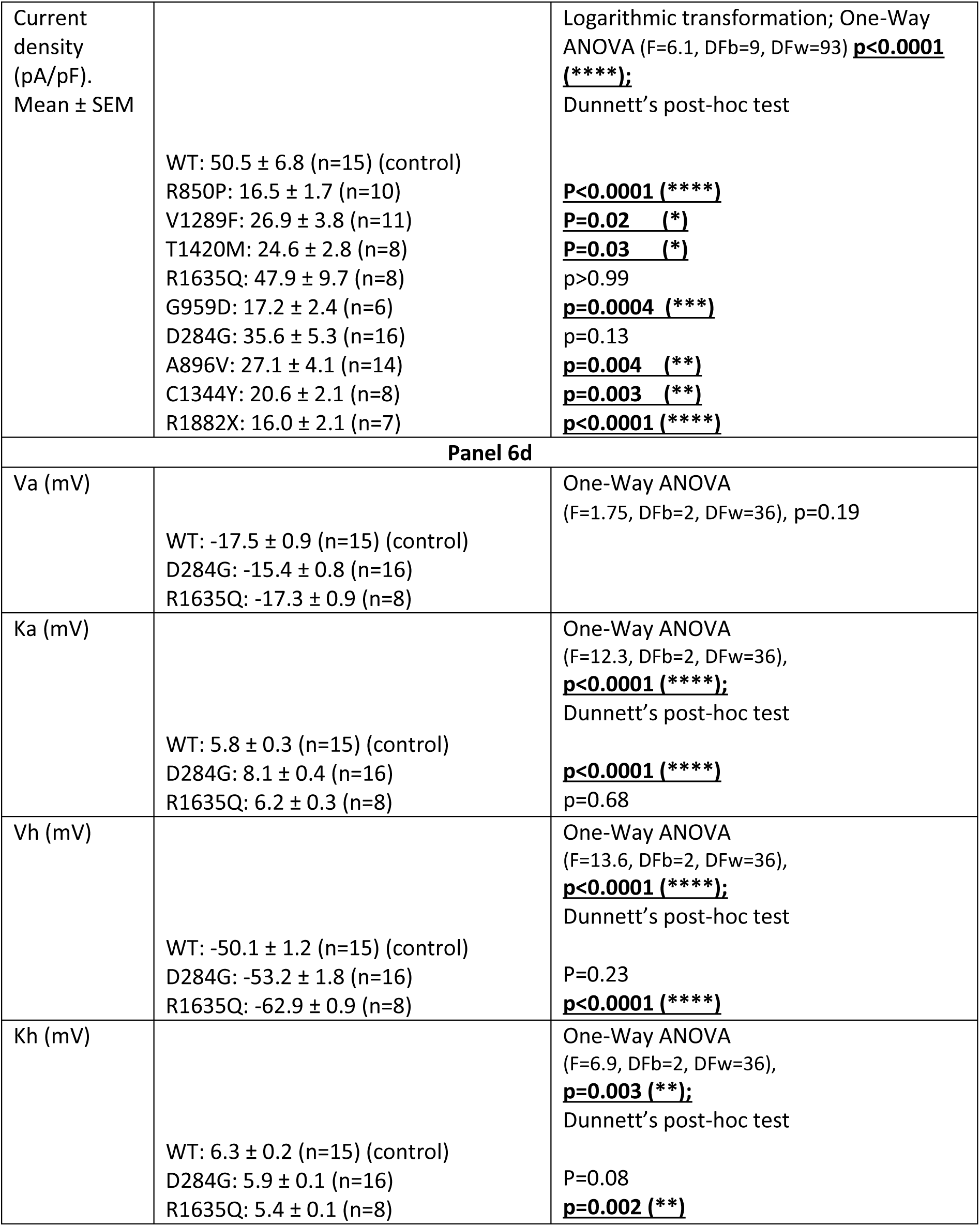
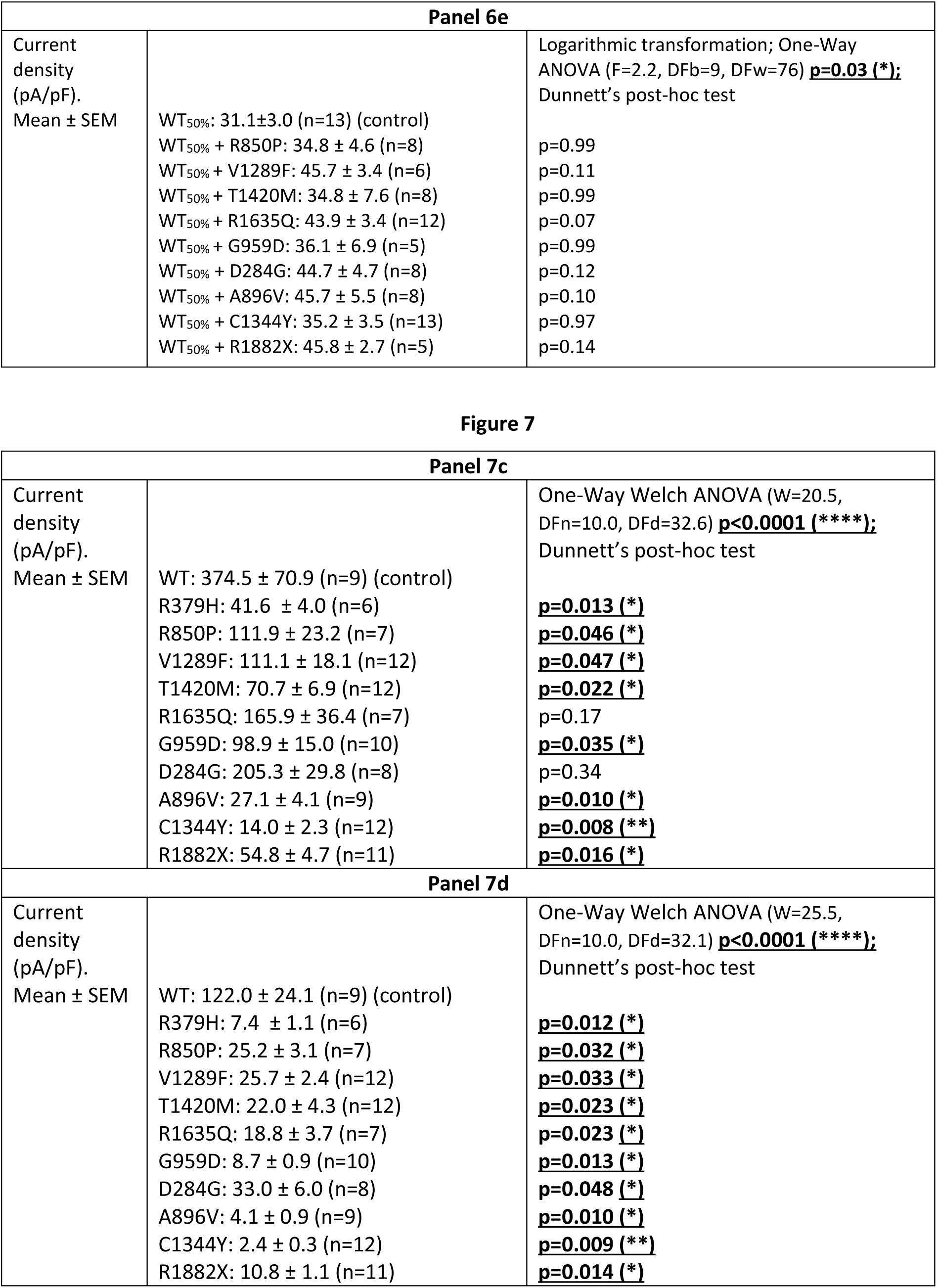

